# Effect of cytochrome CYP2C19 metabolizing activity on antidepressant response and side effects: meta-analysis of data from genome-wide association studies

**DOI:** 10.1101/259838

**Authors:** Chiara Fabbri, Katherine E. Tansey, Roy H. Perlis, Joanna Hauser, Neven Henigsberg, Wolfgang Maier, Ole Mors, Anna Placentino, Marcella Rietschel, Daniel Souery, Gerome Breen, Charles Curtis, Sang-Hyuk Lee, Stephen Newhouse, Hamel Patel, Michael O’Donovan, Glyn Lewis, Gregory Jenkins, Richard M. Weinshilboum, Anne Farmer, Katherine J. Aitchison, Ian Craig, Peter McGuffin, Koen Schruers, Joanna M. Biernacka, Rudolf Uher, Cathryn M. Lewis

**Author notes:** Corresponding author: Prof. Cathryn Lewis, Social, Genetic and Developmental Psychiatry Centre (MRC), Institute of Psychiatry, Psychology and Neuroscience – PO80, De Crespigny Park, Denmark Hill, London, United Kingdom, SE5 8AF.

## Abstract

Cytochrome (CYP) P450 enzymes have a primary role in antidepressant metabolism and variants in these polymorphic genes are targets for pharmacogenetic investigation. This is the first meta-analysis to investigate how *CYP2C19* polymorphisms predict citalopram/escitalopram efficacy and side effects.

*CYP2C19* phenotypes comprise poor metabolizers (PM), intermediate and intermediate+ metabolizers (IM; IM+), extensive and extensive+ metabolizers (EM [wild type]; EM+) and ultra-rapid metabolizers (UM) defined by the two most common *CYP2C19* functional polymorphisms (rs4244285 and rs12248560) in Caucasians. These polymorphisms were genotyped or imputed from genome-wide data in four samples treated with citalopram or escitalopram (GENDEP, STAR*D, GenPod, PGRN-AMPS). Treatment efficacy was percentage symptom improvement and remission. Side effect data were available at weeks 2–4, 6 and 9 in three of the investigated samples. A fixed-effects meta-analysis was performed using EM as the reference group.

Analysis of 2558 patients for efficacy and 2037 patients for side effects showed that PMs had higher symptom improvement (SMD=0.43, CI=0.19–0.66) and higher remission rates (OR=1.55, CI=1.23–1.96) compared to EMs. At weeks 2–4, PMs showed higher risk of gastro-intestinal (OR=1.26, CI=1.08–1.47), neurological (OR=1.28, CI=1.07–1.53) and sexual side effects (OR=1.52, CI=1.23–1.87; week 6 values similar). No difference was seen at week 9 or in total side effect burden. PMs did not have higher risk of dropout at week 4 compared to EMs. Antidepressant dose was not different among *CYP2C19* groups.

*CYP2C19* polymorphisms may provide helpful information for guiding citalopram/escitalopram treatment, despite PMs are relatively rare among Caucasians (~2%).

## 1. Introduction

Major depressive disorder (MDD) is a leading cause of disability-adjusted life years worldwide (GBD 2015 Disease and Injury Incidence and Prevalence Collaborators, 2016). Although anti-depressant drugs can be an effective therapy, remission rates are disappointing, largely as a consequence of high variability in efficacy among individuals combined with early discontinuation or poor compliance due to side effects (Hodgson et al., 2012); (Crawford et al., 2014). Genetic variants are considered key modulators of antidepressant efficacy and side effects (Cacabelos et al., 2012). Common variants were estimated to explain approximately 42% of inter-individual variability in antidepressant response (Tansey et al., 2013), confirming the role of genetic polymorphisms as promising markers to provide personalized treatments.

Previous pharmacogenetic studies for antidepressant efficacy and side effects have focused on genes involved in antidepressant mechanisms of action (pharmacodynamics) or in antidepressant transport/metabolism (pharmacokinetics), including the cytochrome P450 genes (CYP450) (Fabbri and Serretti, 2015). These CYP450 genes are included in commercial pharmacogenetic tests (e.g. GeneSight Psychotropic, Genecept Assay™, YouScript Psychotropic (GTR: Genetic Testing Registry, 2017)). They form promising targets for personalizing antidepressant treatment, since they are responsible for antidepressant drug metabolism and their polymorphisms define phenotypic groups with different level of metabolic activity (Porcelli et al., 2011). An association between CYP450 metabolizer status (CYP450 phenotypes) and metabolite plasma levels has been consistently reported for antidepressants, but the association of CYP450 phenotypes with antidepressant efficacy and side effects is more controversial (Porcelli et al., 2011).

*CYP2C19* is the primary CYP450 isoform responsible for the metabolism of citalopram and escitalopram, two commonly prescribed SSRIs (selective serotonin reuptake inhibitors) (Hicks et al., 2015). Elevated drug concentrations have been observed in *CYP2C19* poor metabolizers (PMs), which may increase the risk of adverse drug reactions, while *CYP2C19* ultrarapid metabolizers (UMs) may have lower exposure to these drugs leading to treatment failure. *CYP2C19-*adjusted doses for citalopram and escitalopram have been estimated, but these were based on observed differences in drug pharmacokinetics, not differences in clinical outcomes of efficacy and side effects (Hicks et al., 2015).

Inconsistent associations between *CYP2C19* phenotypes and citalopram/escitalopram outcomes have been observed, and several factors may have led to the contradictory results (Peters et al., 2008) (Mrazek et al., 2011)(Hodgson et al., 2014)(Hodgson et al., 2015):

1. Only a weak correlation exists between SSRI dose and efficacy and drug plasma levels may not be associated with either efficacy or side effects (Jakubovski et al., 2016) (Hodgson et al., 2014) (Hodgson et al., 2015);
2. Pharmacodynamic mechanisms may modulate the association between *CYP2C19* phenotypes and citalopram/escitalopram efficacy and some side effects, weakening the association between pharmacokinetic parameters and treatment outcomes (Jukić et al., 2016);
3. *CYP2C19* PM phenotypes are rare, and studies may have lacked power to detect a pharmacogenetic association with this phenotype.

In this study, we present the first meta-analysis to investigate association between *CYP2C19* phenotypes and citalopram/escitalopram efficacy and side effects. This large study aimed to identify a link between *CYP2C19* phenotypes and treatment outcomes and to determine whether dose adjustments based on *CYP2C19* phenotypes should be part of personalized medicine for antidepressant treatment.

## 2. Experimental procedures

### 2.1. Samples

#### 2.1.1. GENDEP

The Genome-Based Therapeutic Drugs for Depression (GENDEP) project was a 12-week partially randomized open-label pharmacogenetic study with two active treatment arms. 867 patients with unipolar depression (ICD-10 or DSM-IV criteria) aged 19–72 years were recruited at nine European centres. Eligible participants were allocated to flexible-dosage treatment with either escitalopram (10–30 mg daily) or nortriptyline. Only 499 patients treated with escitalopram were included in the current meta-analysis. Severity of depression was assessed weekly by the Montgomery-Asberg Depression Rating Scale (MADRS) (Montgomery and Asberg, 1979), Hamilton Rating Scale for Depression (HRSD–17) (Hamilton, 1967) and other measures. Side effects were assessed at baseline and then weekly using the Antidepressant Side-Effect Checklist (ASEC) and UKU Side Effect Rating Scale, with good agreement between them. The ASEC data were analysed for this study, since they have lower rates of missing data (Uher et al., 2009). Detailed information about the GENDEP study has been previously reported (Uher et al., 2010).

#### 2.1.2 STAR*D

The Sequenced Treatment Alternatives to Relieve Depression (STAR*D) study was a NIMH-funded study to determine the effectiveness of different treatments for patients with MDD who have not responded to the first antidepressant treatment. Non-psychotic MDD (DSM-IV criteria) patients with age between 18 and 75 years were enrolled from primary care or psychiatric outpatient clinics. Severity of depression was assessed using the 16-item Quick Inventory of Depressive Symptomatology-Clinician Rated (QIDS-C16) (Trivedi et al., 2004) at baseline, weeks 2, 4, 6, 9, and 12. Side effects were measured at the same time points using the Patient-Rated Inventory of Side Effects (PRISE). This study uses data from level 1, where all patients received citalopram. Detailed description of the study design and population are reported elsewhere (Rush et al., 2004).

#### 2.1.3. PGRN-AMPS

The Pharmacogenomic Research Network Antidepressant Medication Pharmacogenomic Study (PGRN-AMPS) included 529 participants with nonpsychotic MDD recruited at inpatient and outpatient practices of the Department of Psychiatry and Psychology, Mayo Clinic, Rochester, Minnesota. Participants were offered an eight-week course of treatment with either citalopram or escitalopram and depressive symptoms were rated using QIDS-C16 as in STAR*D. Side effects were assessed using the PRISE scale at weeks 4 and 8. Further details were reported elsewhere (Ji et al., 2013).

#### 2.1.4. GenPod

The GENetic and clinical Predictors Of treatment response in Depression (GenPod) was a multi-centre randomized clinical trial of 601 patients recruited in primary care who had an ICD-10 diagnosis of major depression of at least moderate severity as assessed by the Clinical Interview Schedule-Revised (CIS-R) (Lewis et al., 1992) and the Beck Depression Inventory (BDI) (Beck et al., 1961). Individuals were randomly allocated to either reboxetine (4 mg twice daily) or citalopram (20 mg/day). 240 patients of European ancestry and treated with citalopram were included in this meta-analysis. Further details about this study can be found elsewhere (Thomas et al., 2008).

### 2.2. Outcomes

#### 2.2.1. Treatment efficacy

Treatment efficacy was measured by percentage symptom improvement and by remission at study endpoint. Continuous measures, such as percentage improvement, capture more information and have higher power than cutoff-based dichotomous measures, however remission has a particular clinical relevance since it is associated with MDD prognosis (Streiner, 2002)(Gaynes et al., 2009). The percentage symptom improvement was corrected for possible confounding variables (age, baseline severity, and center for multi-center studies) and then standardized to allow comparability across studies.

Remission was defined as a binary variable according to standard definitions (HRSD–17 ≤ 7 in GENDEP; QIDS-C16 ≤ 5 in STAR*D and PGRN-AMPS; BDI < 10 in GenPod). In GENDEP symptom improvement was calculated using the MADRS scale similarly to previous studies (Uher et al., 2010) while HRSD–17 was used to define remission given the stronger consensus about the threshold to identify remission on this scale in contrast to MADRS, where different definitions of remission have been reported (Li et al., 2016) (Jacobsen et al., 2015).

HRSD–17 and QIDS-C16 missing values at follow-up were imputed using the best unbiased estimate from a mixed-effect linear regression model, with fixed linear and quadratic effects of time and random effects of individual and center of recruitment, following previously reported methods (Uher et al., 2010).

#### 2.2.2. Side effects

Measures of side effects were available in GENDEP, STAR*D and PGRN-AMPS. In GENDEP we chose to use the ASEC because data was more complete than the UKU (Uher et al., 2009). In STAR*D and PGRN-AMPS side effects were assessed using the PRISE scale. Both scales use a rating of severity for each side effect (coded 0–3 in ASEC, and 0–2 in PRISE) which was dichotomized (0=absent, 1=present) for the meta-analysis. Side effects were grouped in categories that were assessed in both samples: gastro-intestinal (dry mouth, diarrhea, constipation, nausea or vomiting), cardiovascular (palpitations, dizziness or feeling light-headed on standing), central nervous system (headache, tremor, feeling like the room is spinning), sleep (insomnia, drowsiness or oversleeping) and sexual (loss of desire, trouble achieving orgasm, trouble with erection). These categories were analysed as dichotomous variables (presence of at least one side effect in each category). To assess the overall severity of side effects across both studies, we summed the number of side effects reported, and dichotomized at the 3^rd^ quartile of the distribution in each sample. Study retention at week 4 was compared among *CYP2C19* phenotypes since patients who did not benefit from treatment or had troubling side effects are expected to be lost from follow-up early in the study.

Antidepressant-induced side effects are more frequent at the beginning of treatment and then decrease (Uher et al., 2009). We therefore meta-analysed side effects at weeks 2–4 (no assessment was performed at week 2 in PGRN-AMPS), week 6 and weeks 8–9 (no assessment was performed at week 8 in STAR*D while in GENDEP we used week 8 data because of lower missing rate compared to week 9).

In GENDEP side effects were common at baseline in medication-free patients (Uher et al., 2009). We therefore performed a sensitivity analysis excluding side effects there were present also at baseline in drug-free GENDEP patients.

### 2.3. Genotyping and definition of *CYP2C19* phenotypes

*CYP2C19* phenotypes comprise poor metabolizers (PM), intermediate and intermediate+ metabolizers (IM; IM+), extensive and extensive+ metabolizers (EM [wild type]; EM+) and ultra-rapid metabolizers (UM) defined by the two most common *CYP2C19* functional polymorphisms (rs4244285 and rs12248560) which capture the CYP2C19 *1, *2 and *17 functional alleles (Supplementary Table 1) (Hodgson et al., 2014). These polymorphisms were directly genotyped in GENDEP using the AmpliChip CYP450 test (Hodgson et al., 2014) and they were imputed in the other samples using the Haplotype Reference Consortium (HRC version r1.1 2016) panel as reference and Minimac3. Pre-imputation quality control was performed according to standard criteria (variants with missing rate ≥ 5%; monomorphic variants; subjects with genotyping rate < 97%; subjects with gender discrepancies; subjects with abnormal heterozygosity; related subjects (identity by descent (IBD) >0.1875 (Anderson et al., 2010)); population outliers according to Eigensoft analysis of linkage-disequilibrium-pruned genetic data (Price et al., 2006); and non-white subjects). Imputation quality was assessed using R^2^ (Li et al., 2010) and comparing imputed and genotyped *CYP2C19* phenotypes in GENDEP.

### 2.4. Statistical analysis

Individual-level phenotypes and genotypes were available for all studies. A fixed-effects meta-analysis was performed with the R package “Netmeta” (https://cran.r-project.org/web/packages/netmeta/index.html). This package has been created for performing network meta-analysis and it was useful for this study since multiple groups needed to be compared to the reference group even if there were not indirect comparisons (i.e. all the studies provided data for each of the considered *CYP2C19* phenotypes). Phenotypic groups were compared using the wild-type EM as the reference group. A random-effects meta-analysis was carried out for completeness and comparison of findings. Standardized mean difference (SMD) or odds ratio (OR) with 95% confidence intervals (CI) were calculated. Heterogeneity across studies was assessed using I^2^ and Cochran’s Q (Higgins et al., 2003).

This meta-analysis provided 80% power to identify an effect size (SMD) of *d*=0.40 when comparing PMs (the smallest group, n=51) with EMs (the reference group, n=1049) for a continuous outcome and OR=2.21 for a binary outcome, at a significance level of 0.05 (Faul et al., 2007).

We estimated that a corrected p value of 0.008 would account for the six independent tests that were carried out (improvement and response were correlated and considered as one test; gastrointestinal side effects, cardiovascular side effects, sleep side effects, sexual side effects, and CNS side effects were considered as independent outcomes). Side effects at different weeks are not independent and *CYP2C19* metabolic groups are not considered independent (they all derive from two functional SNPs in the gene), and they have specific functional meaning.

## 3. Results

A description of the clinical-demographic characteristics of the included samples is provided in Supplementary Table 2. There was no difference in mean citalopram or escitalopram dose by *CYP2C19* phenotypes at study endpoint in GENDEP, STAR*D and PGRN-AMPS (dose information was not available in GenPod). The distribution of phenotypic groups in the analysed samples is reported in Supplementary Table 3A. Imputation quality was high in all samples for both polymorphisms (R^2^ between 0.95 and 0.99 (Li et al., 2010)). GENDEP participants had 97.6% consistency between genotyped and imputed SNPs (Supplementary Table 3B).

### 3.1. Treatment efficacy

In total, 2558 patients were included in the meta-analysis. The distribution of efficacy outcomes across *CYP2C19* phenotypes was reported in Supplementary Table 4. Compared to EMs, PMs had higher symptom improvement scores (SMD=0.43, CI=0.19–0.66, p=0.00037) and higher remission rates (OR=1.55, CI=1.23–1.96, p=0.00025), with low or absent heterogeneity (I^2^ was 11.5% and 0%, respectively). Other *CYP2C19* phenotypes did not show different outcomes compared to EMs (Figure 1). Results did not change using a random-effects model.

**Figure 1:**
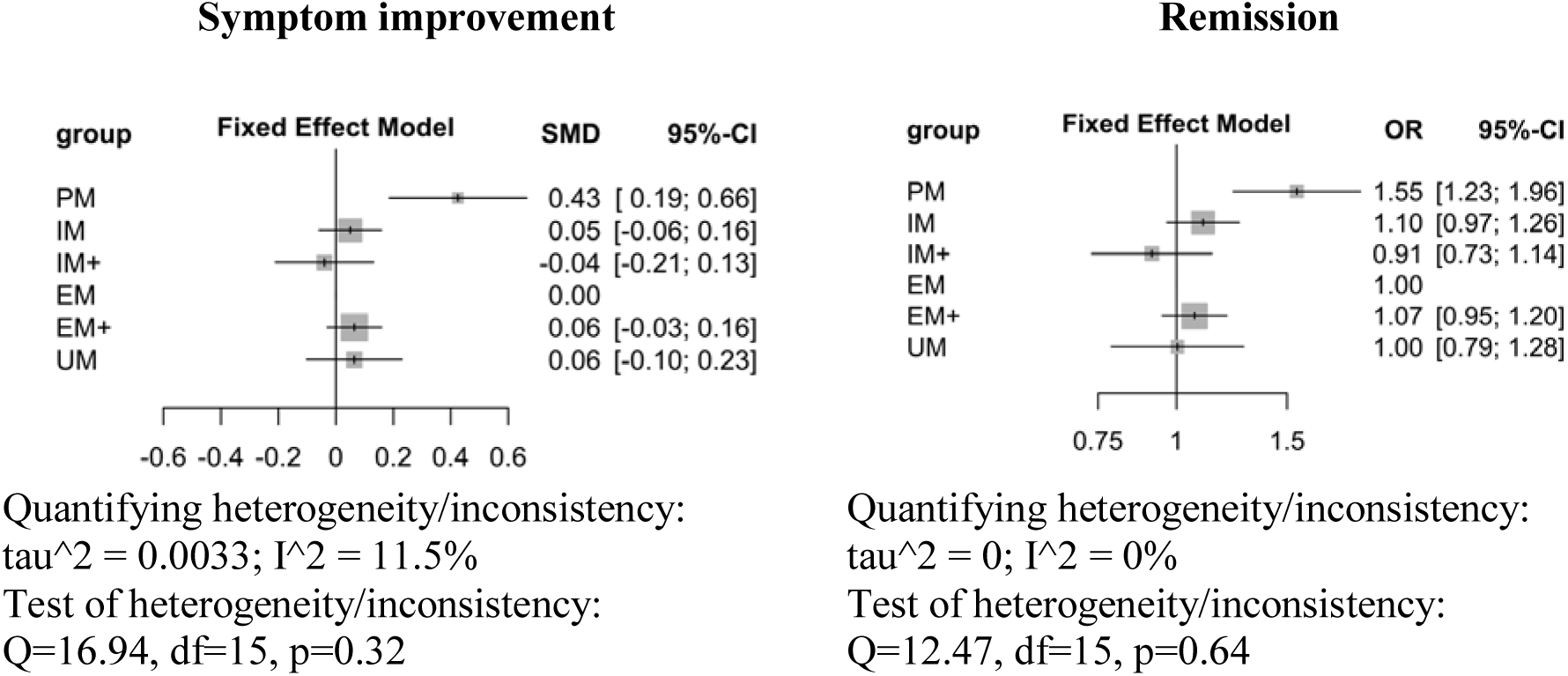
meta-analysis results for improvement and remission. PM=poor metabolizers: IM=intermediate metabolizers; IM+= intermediate metabolizers plus; EM=extensive metabolizers; EM+= extensive metabolizers+; UM=ultrarapid metabolizers. EM was taken as reference group. SMD=standardized mean difference. CI=confidence interval.

### 3.2. Treatment side effects

Across STAR*D, GENDEP and PGRN-AMPS 2037 patients were included in the analysis. The distribution of side effects across *CYP2C19* phenotypes was reported in Supplementary Table 5. At weeks 2–4, PMs showed higher risk of gastro-intestinal side effects (OR=1.26, CI=1.08–1.47, p=0.0033), of CNS side effects (OR=1.28, CI=1.07–1.53, p=0.0068) and of sexual side effects (OR=1.52, CI=1.23–1.87, p=0.0001) (Figure 2). Considering a corrected p threshold of 0.008, all these side effects were significantly more frequent in PMs. At week 6, PMs showed higher risk of sexual side effects (OR=1.64, CI=1.23–2.17, p=0.0007) but no higher risk of other side effects. For all these comparisons heterogeneity was low (I^2^ range 0%-24%). No difference was seen at week 8–9 for any side effect, except a weak non-significant trend for sexual side effects; no difference in total side effects burden was observed at any time point (Figure 2). *CYP2C19* IM+ group was the only phenotype to show higher risk of drop out at week 4 (OR=1.80, 95% CI=1.08–3.00, p=0.024), but this association did not survive multiple-testing correction. PMs did not show higher risk of dropout at week 4 (OR=1.16, CI=0.38–3.58). Other *CYP2C19* phenotypic groups did not show relevant differences compared to EMs, except lower risk of cardiovascular side effects and sleep side effects in EM+ at weeks 2–4 (OR=0.77, CI=0.64–0.92, p=0.0048) and 6 (OR=0.84, CI=0.75–0.95, p=0.0039), respectively, and higher risk of CNS side effects at week 8 in UMs (OR=1.26, 95% CI=1.04–1.53, p=0.019), but the latter did not survive multiple-testing correction.

The use of a random-effects model did not change the results.

**Figure 2:**
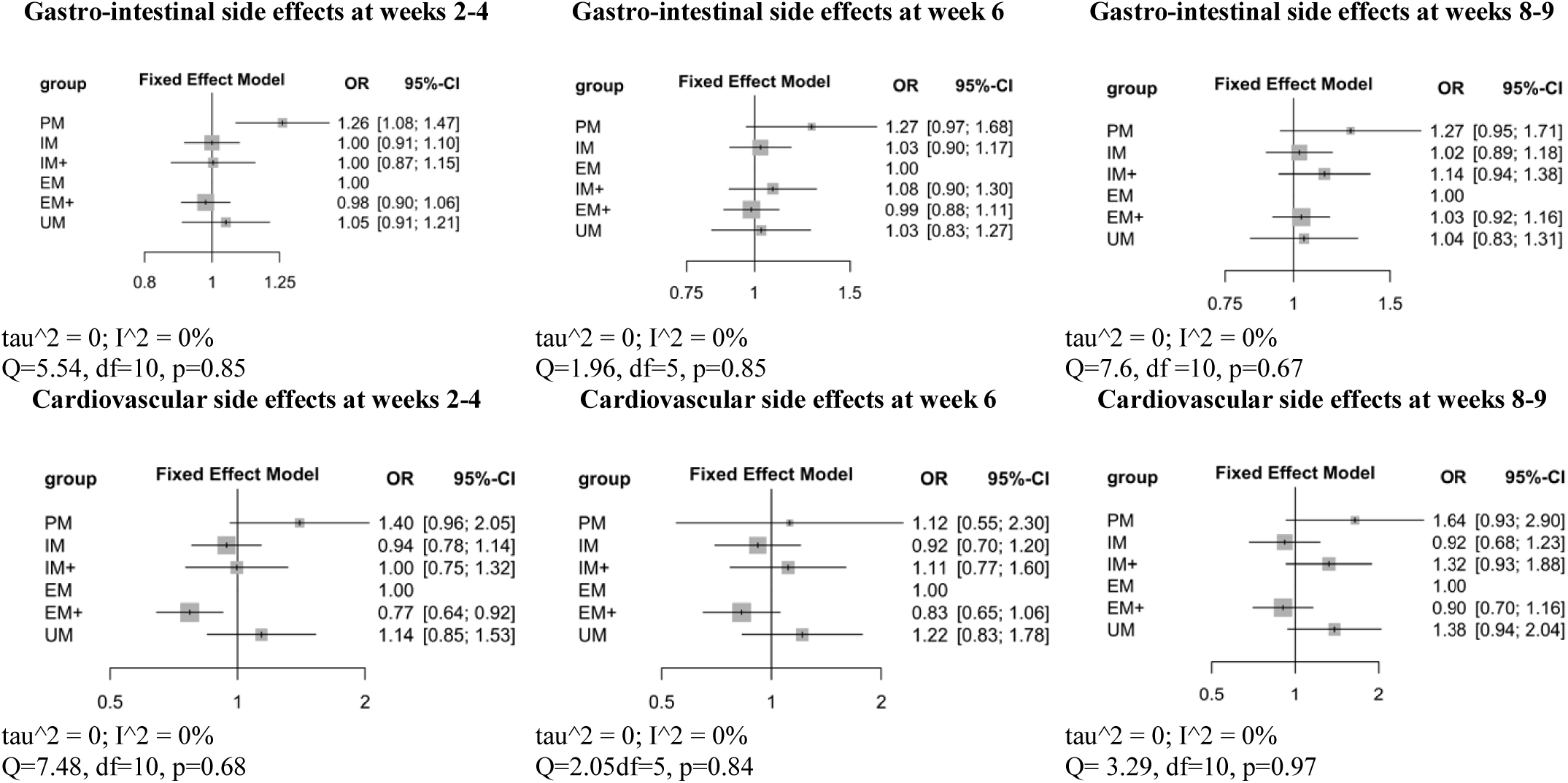
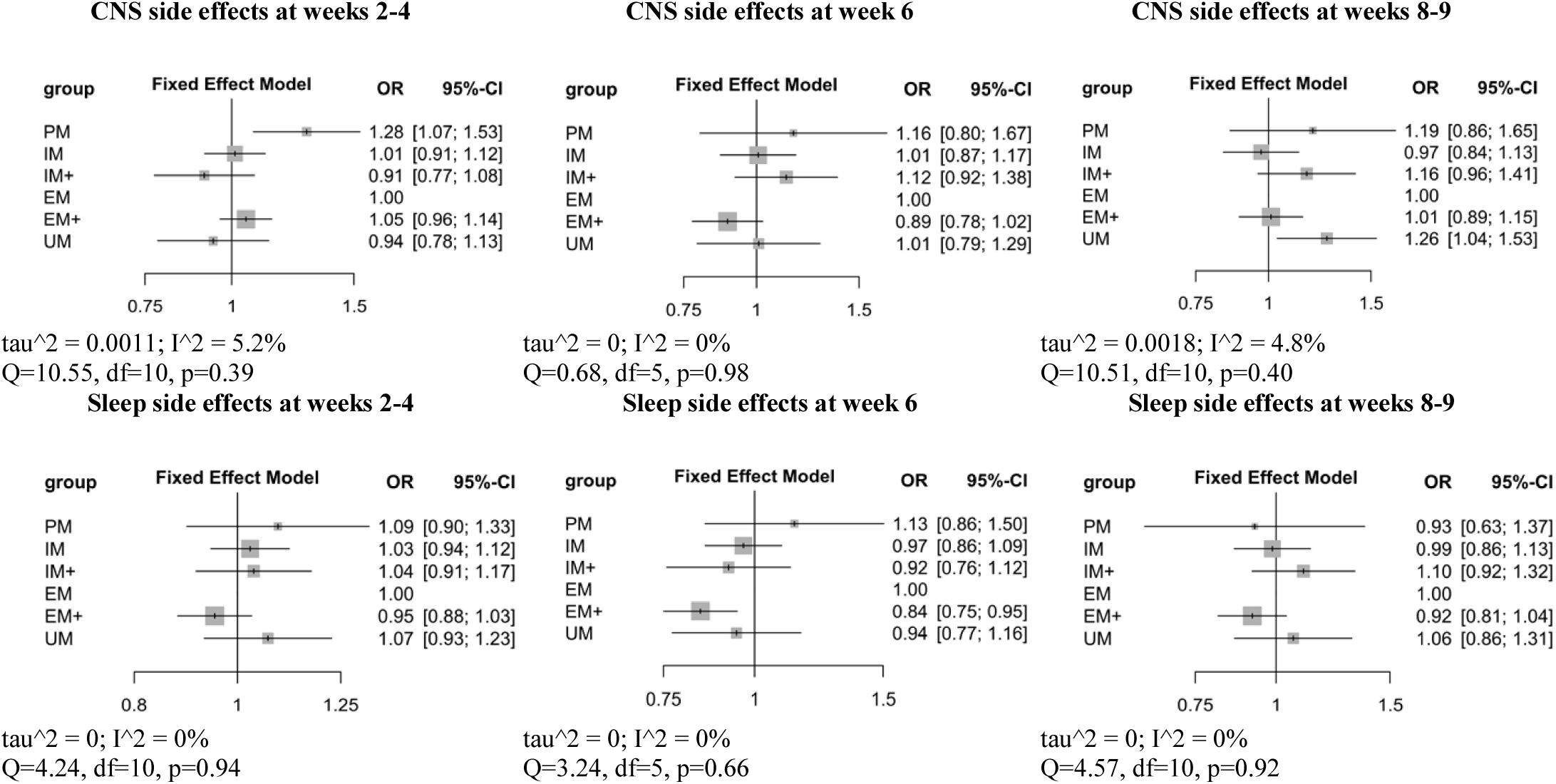
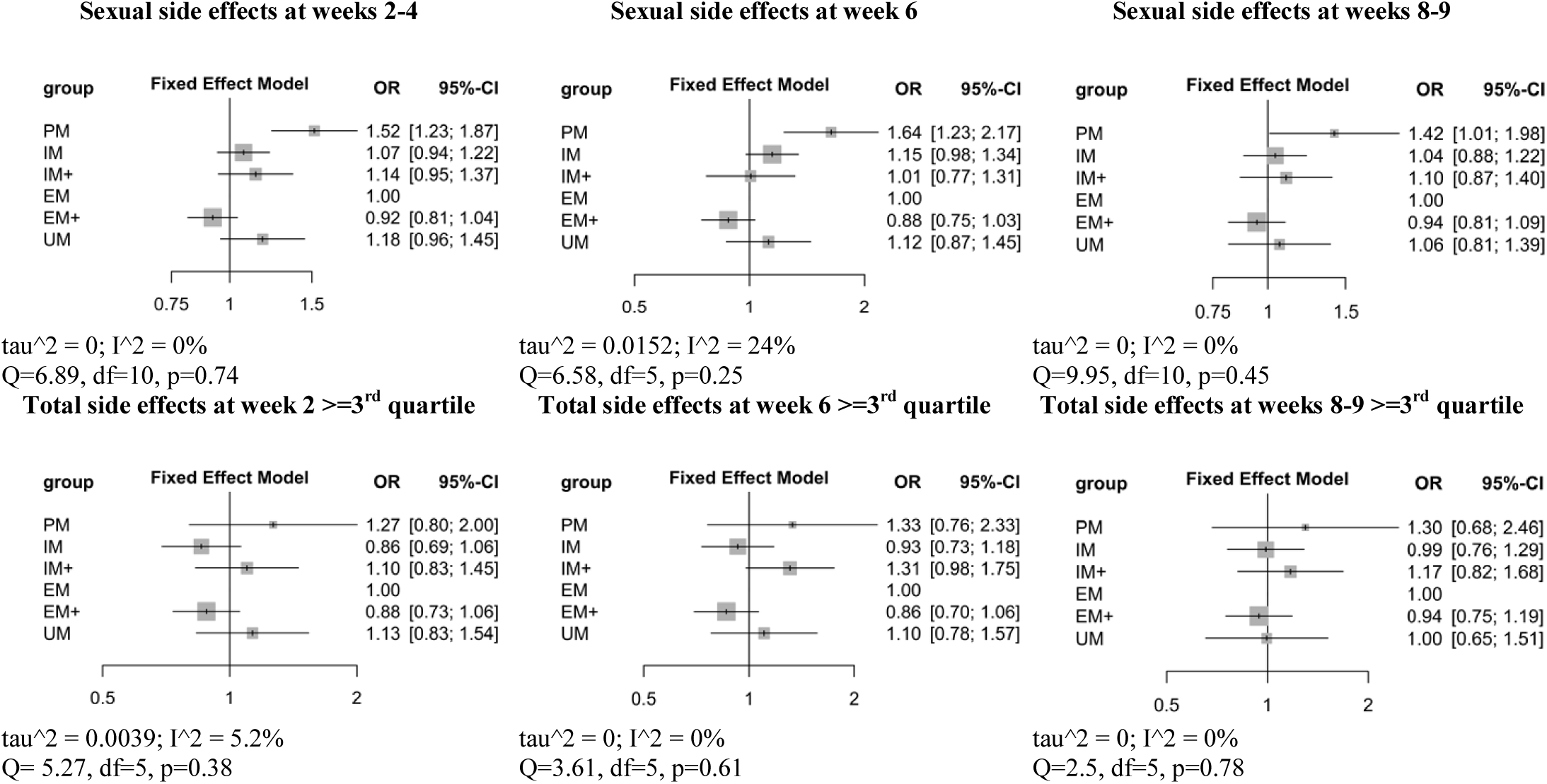
meta-analysis results for side effects. PM=poor metabolizers: IM=intermediate metabolizers; IM+= intermediate metabolizers plus; EM=extensive metabolizers; EM+= extensive metabolizers+; UM=ultrarapid metabolizers. EM was taken as reference group. SMD=standardized mean difference. CI=confidence interval. For each comparison heterogeneity is quantified using tau^2, I^2 and assessed using Q test.

Excluding those side effects there were already present at baseline in drug-free patients in GENDEP, results did not change, except that PMs showed higher risk of gastro-intestinal side effects also at week 6 (OR=1.47, CI=1.13–1.92, p=0.004). In addition, the trend of higher sexual side effects in PMs at weeks 8–9 was not observed, the lower risk of cardiovascular side effects in EM+ at weeks 2–4 became a non-significant trend (OR=0.82, CI=0.67–0.99) and there was a nonsignificant trend of higher gastro-intestinal side effects in PMs at weeks 8–9 (OR=1.35, CI=1.01–1.81).

## 4. Discussion

This study shows that *CYP2C19* PMs had higher symptom improvement and higher remission probability compared to EMs during treatment with citalopram or escitalopram (Figure 1). The observed SMD of 0.43 in symptom improvement between PMs and EMs is statistically considered close to a medium effect size (0.50) (Faraone, 2008). Statistical outcomes cannot be equated with clinical relevance and a clinical relevance cutoff of SMD=0.24 was proposed based on the effect size observed for antidepressant drugs (SMD=0.31, CI=0.27–0.35) and psychotherapy (SMD=0.25, CI= 0.14–0.36) in depression (Cuijpers et al., 2014). Other *CYP2C19* phenotypes, including UMs, showed no differences in efficacy outcomes compared to EMs. In addition to increased treatment efficacy, PMs showed higher risk of gastro-intestinal, CNS and sexual side effects early in treatment (particularly during the first 2–4 weeks), but not later in treatment (weeks 8–9) (Figure 2). At week 4, PMs did not show a higher burden of total side effects and had no higher risk of dropout. Mean antidepressant dose was not different among *CYP2C19* metabolizing groups. These results suggest that although some side effects were more common in PMs in the first weeks of treatment, overall they were not more troubling than in other *CYP2C19* groups and they may be balanced by higher improvement in depressive symptoms.

These findings are consistent with a previous STAR*D study that investigated remission and tolerance to citalopram (Mrazek et al., 2011), where tolerance represents a measure of side effect level. Tolerance was defined as continuation of citalopram treatment after the completion of Level 1 of the STAR*D trial. Previous studies in GENDEP and STAR*D failed to establish association between *CYP2C19* metabolizer status (PM vs. EM) and response, side effects or study retention (Peters et al., 2008) (Hodgson et al., 2015)(Hodgson et al., 2014), but individual studies would have limited power given the low number of subjects with PM phenotype (~2% of all patients analysed), particularly in GENDEP which has only six PM subjects. A previous analysis of *CYP2C19* in GENDEP used different definitions of side-effect, investigating each ASEC item and the sum of ASEC items (Hodgson et al., 2015).

No difference in treatment efficacy or side effects was identified between UMs and EMs, except for a non-significant higher risk of CNS side effects only at weeks 8–9 (Figure 2) that was probably the effect of random noise.

The only phenotypic group that showed lower risk of side effects was EM+ (lower risk of sleep side effects at week 6 and of cardiovascular side effects at weeks 2–4), suggesting that weak differences may depend on metabolic level but the UM group may have not provided enough power to observe them (~4–5% of patients were UMs in the analysed samples).

In addition to pharmacokinetic mechanisms, pharmacodynamic mechanisms may be involved in the association between *CYP2C19* and antidepressant response, since *CYP2C19* activity was reported to influence central neurotransmitters and neurotrophins relevant to antidepressant mechanisms of action (Jukić et al., 2016).

Our results conflict with the recommendation, based on pharmacokinetic parameters, of a 50% reduction in the starting dose of citalopram/escitalopram in *CYP2C19* PMs (Hicks et al., 2015), since we showed that a standard dose was associated with greater efficacy without higher drop-out rates or higher total burden of side effects. Antidepressant treatment with citalopram/escitalopram may be particularly indicated in *CYP2C19* PMs given the efficacy profile, if appropriate clinical support and monitoring is provided and the patient is informed of potential side effects at the beginning of the treatment. Effective plasma (and brain) drug concentrations may be reached in a higher proportion of PMs than other phenotypes, at the price of more frequent early side effects. The good tolerability profile of citalopram/escitalopram implies that these side effects are usually not troubling, which may not be true for other antidepressants, such as tricyclic antidepressants (TCAs) or venlafaxine (Cipriani et al., 2012)(Cipriani et al., 2009). It should be noted that TCAs and venlafaxine have specific profiles of efficacy and they represent valid alternatives to SSRIs as currently reported in clinical guidelines, but it should not be assumed that the current results referred to *CYP2C19* PMs can be applied to antidepressants different from citalopram and escitalopram.

The limitations and strengths of this study should be considered. This was the first meta-analysis to investigate the role of *CYP2C19* phenotypes in citalopram/escitalopram efficacy/side effects, individual level data were available in all samples and the total sample size was the largest ever used for investigating this topic. On the other hand, PMs are rare in the Caucasian population resulting in limited power to identify differences involving this group even in this sample of 2558 patients. Side effect assessment was not available in all samples, and at weeks 6 and 8–9 part of patients dropped from the study and side effects data could not be imputed because it would be unreliable. At weeks 6 and 8–9, respectively, side effects were available in 84.4% and 73.6% of the initial sample in STAR*D, while in 85.9% and 83.3% of the initial sample in GENDEP. In PGRN-AMPS 0.87% of patients initially included had side effect data at week 4 and 80% at week 8. Our findings suggest that *CYP2C19* PMs may benefit from standard doses of citalopram/escitalopram, with a higher response than other phenotypes. No conclusions could be drawn for UMs since which showed no significant differences in outcomes compared to EMs, and the study was probably under-powered to detect weak effects. EM+ was the only group showing lower risk of some side effects compared to EMs. We observed no to low heterogeneity among studies for both efficacy and side effects. For the former group all samples showed similar better outcome in PMs compared to EMs except GENPOD, which included only three PM patients explaining the marginal effect on heterogeneity. Finally, the possible confounding effect of *CYP2C19* enhancers/inhibitors was not assessed, but a previous analysis in GENDEP concluded that the exclusion of subjects with concomitant use of enhancers/inhibitors did not change the pattern of results (Hodgson et al., 2014). In conclusion, this meta-analysis shows good efficacy in *CYP2C19* poor metabolisers with citalopram/escitalopram, contrasting previous pharmacokinetic findings (Hicks et al., 2015). Our results show better treatment outcomes in PMs treated with standard doses with no relevant impact on late side effects (after the 6^th^ week of treatment). Careful information for patients and monitoring of side effects during the early phase of treatment are recommended. Other *CYP2C19* phenotypes, including UMs, did not show differences in efficacy or side-effect outcomes compared to EMs. An interesting implication of this study is the possibility to derive *CYP2C19* metabolic groups from standard genome-wide data with a good level of quality.

## Acknowledgments/role of the funding source

This report represents independent research part-funded by the National Institute for Health Research (NIHR) Biomedical Research Centre at South London and Maudsley NHS Foundation Trust and King’s College London. The views expressed are those of the authors and not necessarily those of the NHS, the NIHR, or the Department of Health.

We thank the NIMH for providing access to data on the STAR-D sample. We also thank the authors of previous publications in this dataset, and foremost, we thank the patients and their families who agreed to be enrolled in the study. Data and biomaterials were obtained from the limited access datasets distributed from the NIH-supported ‘‘Sequenced Treatment Alternatives to Relieve Depression’’ (STAR*D). The study was supported by NIMH Contract No. N01MH90003 to the University of Texas Southwestern Medical Center. The ClinicalTrials.gov identifier is NCT00021528.

The GENDEP project was supported by a European Commission Framework 6 grant (contract reference: LSHB-CT-2003-503428). The Medical Research Council, United Kingdom, and GlaxoSmithKline (G0701420) provided support for genotyping.

The NEWMEDS study was funded by the Innovative Medicine Initiative Joint Undertaking (IMI-JU) under grant agreement n° 115008 of which resources are composed of European Union and the European Federation of Pharmaceutical Industries and Associations (EFPIA) in-kind contribution and financial contribution from the European Union’s Seventh Framework Programme (FP7/2007-2013). EFPIA members Pfizer, Glaxo Smith Kline, and F. Hoffmann La-Roche have contributed work and samples to the project presented here. The funders had no role in study design, data collection and analysis, decision to publish, or preparation of the manuscript.

The PGRN-AMPS dataset used for the analyses described in this manuscript was obtained from dbGaP (study accession phs000670.v1.p1). PGRN-AMPS was supported, in part, by NIH grants RO1 GM28157, U19 GM61388 (The Pharmacogenomics Research Network), U01 HG005137, R01 CA138461, P20 1P20AA017830-01 (The Mayo Clinic Center for Individualized Treatment of Alcohol Dependence), and a PhRMA Foundation Center of Excellence in Clinical Pharmacology Award.

Dr Rudolf Uher is supported by the Canada Research Chairs Program.

